# Parallel genetic adaptation of *Bacillus subtilis* to different plant species

**DOI:** 10.1101/2023.03.17.533125

**Authors:** Guohai Hu, Yue Wang, Christopher Blake, Mathilde Nordgaard, Xin Liu, Bo Wang, Ákos T. Kovács

## Abstract

Plant growth-promoting rhizobacteria benefit plants by stimulating their growth or protecting them against phytopathogens. Rhizobacteria must colonise and persist on plant roots to exert their benefits. However, little is known regarding the processes by which rhizobacteria adapt to different plant species, or behave under alternating host plant regimes. Here, we used experimental evolution and whole-population whole-genome sequencing to analyse how *Bacillus subtilis* evolves on *Arabidopsis thaliana* and tomato seedlings, and under an alternating host plant regime, in a static hydroponic setup. We observed parallel evolution across multiple levels of biological organisation in all conditions, which was greatest for the two heterogeneous, multi-resource spatially-structured environments at the genetic level. Species-specific adaptation at the genetic level was also observed, possibly caused by the selection stress imposed by different host plants. Furthermore, a trade-off between motility and biofilm development was supported by mutational changes in motility– and biofilm-related genes. Finally, we identified several condition-specific and common targeted genes in different environments by comparing three different *B. subtilis* biofilm adaptation settings. The results demonstrate a common evolutionary pattern when *B. subtilis* is adapting to the plant rhizosphere in similar conditions, and reveal differences in genetic mechanisms between different host plants. These findings will likely support strain improvements for sustainable agriculture.

**Data summary:** Sequencing data associated with this article are available in the CNGB Sequence Archive (CNSA) [1] of the China National GeneBank DataBase (CNGBdb) [2] under accession numbers CNP0002416 and CNP0003952. Strain data for the DK1042 ancestor are available under accession number CNP0002416.

**Impact statement:** For rhizobacteria to benefit plant growth and protect against phytopathogens, bacteria must colonise and persist on plant roots. Understanding how rhizobacteria adapt to different plant species will assist strain development in sustainable agriculture. To explore the rhizobacterial adaptation process for different plant species and alternating host regimes, *B. subtilis* was experimentally evolved on *A. thaliana* or tomato roots, or an alternating host regime. Both parallel and species-specific adaptation was revealed at the genetic level. Analysis of the trade-off between motility and biofilm formation revealed several condition-specific and commonly targeted genes based on experimentally evolving *B. subtilis* biofilms.

## Introduction

In natural environments, plant growth and yield depend on a plethora of interactions within the complex and dynamic communities of bacteria and fungi [3]. The rhizosphere provides a niche for the mutualistic relationships between plants and microorganisms in the surrounding soil. Plants exude up to 20% of fixed carbon and 15% of nitrogen into their environment [4, 5], and these exudates shape the microbiome. Importantly, the amount and composition of exudates vary depending on plant species, stage of development and environmental stresses [6]. Exudates contribute to rhizobiome formation by recruiting plant growth-promoting rhizobacteria (PGPR) and serving as nutrients [7], and PGPR benefit plant growth and enhance plant tolerance to different abiotic and biotic stresses [8]. *Bacillus subtilis* is a soil-dwelling, biofilm-forming Gram-positive PGPR commonly found in association with plants and their rhizosphere [9]. This bacterial species aids plants in several direct and indirect ways [10]. Some *B. subtilis* strains have been applied as biocontrol agents within agriculture, but the effects vary immensely under field conditions [11, 12]. To exert their beneficial effects, PGPR must colonise and persist on the root surface, which mainly depends on their ability to form biofilms.

Each plant species has a distinct effect on its rhizobiome and this is partly driven by plant-secreted exudates. Root exudates not only provide a nutritional source for soil microorganisms, but also serve as a signal to attract or repel specific groups of microbes [13]. For example, diverse sugars and organic acids induce biofilm formation of *B. subtilis in vitro* or directly on the plant via different molecular mechanisms. Spo0A is a central transcriptional regulator in *B. subtilis* involved in biofilm development [14]. Biofilm formation is triggered when moderate levels of phosphorylated Spo0A (Spo0A-P) are present within a cell. The level of Spo0A-P is controlled by five histidine kinases (KinA−E), which respond to different environmental signals [15]. Chen and colleagues found that *B. subtilis* biofilm formation on tomato roots is dependent on KinD and small signalling molecules, possibly L-malic acid, released by the roots to stimulate *B. subtilis* biofilm *in vitro* [16]. Unlike in tomato, deletion of *kinCD* has only a moderate effect on biofilm formation on *Arabidopsis thaliana* roots [17], suggesting that the kinase requirement of *B. subtilis* for root colonisation varies between plant species. In the case of *A. thaliana*, biofilm formation by *B. subtilis* is triggered by plant polysaccharides pectin, arabinogalactan and xylan, which act both as environmental cues and substrates for matrix synthesis [17]. Moreover, the disaccharide sucrose that is abundantly secreted from plant roots activates a signalling cascade to trigger solid surface motility and promote rhizosphere colonisation by *B. subtilis* [18].

Interestingly, a *B. subtilis* strain isolated from banana rhizosphere and a *Bacillus amylolicefaciens* strain isolated from cucumber rhizosphere colonise their original plant hosts more efficiently than the non-host plant [19], demonstrating species-specific colonisation. Additionally, organic acids detected exclusively in one plant species induce chemotaxis and biofilm formation of the corresponding bacterial isolate from that given host [19]. Chemotaxis and swarming motility are important for plant colonisation [19, 20]; when defective in either, the ability of *B. subtilis* to efficiently colonise tomato seedlings was reduced in a gnotobiotic system [20]. Similarly, mutants lacking flagellum production were unable to colonise the roots of *A. thaliana* in a static hydroponic setup and tomato in a gnotobiotic system [20, 21].

In addition to testing specific mutants, experimental evolution provides a powerful tool for revealing the molecular mechanisms related to plant interactions because microbes adapt to the plant root niche [22]. Experimentally evolved clones of *Bacillus thuringiensis* from *A. thaliana* roots displayed improved root colonisation ability and enhanced swarming, but impaired swimming, in addition to altered bacterial differentiation and pathogenicity [23]. In a similar setup, when *B. subtilis* was adapted to *A. thaliana* roots, evolved isolates demonstrated elevated root colonisation and impaired swimming and swarming motility [24], indicating a possible biofilm-motility trade-off. *Pseudomonas protegens*, adapted to *A. thaliana* rhizosphere under axenic sandy conditions, exhibited enhanced swimming motility and impaired swarming motility [25]. Although these results highlight the existence of an evolutionary trade-off between biofilm development and motility, it remains largely unknown how such mechanisms supervene from an evolutionary perspective.

Since crop rotation regime has great influence on soil multifunctionality and bacterial community in agroecosystems [26], we wondered whether a plant-colonising bacterium adapts differently in an alternating host regime compared with using the same plant throughout the experiment. In addition, we questioned what portion of bacterial adaptation on a plant root will be host-specific. To explore these questions, we employed experimental evolution combined with whole-population metagenome sequencing to analyse how *B. subtilis* evolves on *A. thaliana* and tomato seedlings, as well as in an alternating host regime of these two species in a static hydroponic setup. We observed parallel evolution across multiple levels of biological organisation and a higher gene-level parallelism in populations evolved in an alternating host regime. We also observed species-specific adaptation at the genetic level, which was potentially provoked by specific host plant-imposed selection, either due to root exudates, plant polysaccharides, or certain stress conditions. Additionally, motility-biofilm trade-off was revealed in the mutational landscape of related genes, in addition to reduced swimming and swarming motility. Finally, we identified several condition-specific and shared mutated genes of *B. subtilis* when evolved in different biofilm environments.

## Methods

### Bacterial strains and experimental methods

We used identical materials and methods for tomato and two host-cycling evolution setups as described previously for the *A. thaliana* selection approach [27], including the bacterial strain and culture conditions, seedling preparation, experimental evolution, and morphological diversification experiments.

### Experimental evolution

To initiate experimental evolution we used *A. thaliana* Col-0 seedlings at 14−16 days old and tomato Col-0 seedlings at 8−10 days old. Seedlings were placed in 100 ml reagent bottles containing 27 ml of MSNg and inoculated with 3 ml of *B. subtilis* DK1042 bacterial culture adjusted to OD_600_ = 0.2, resulting in a starting OD_600_ of 0.02. To ensure that roots but not sprouts or leaves provided surface for colonisation, seedlings were pulled through a sterile nylon mesh floating on top of the liquid medium, leaving only the roots submerged in the medium. Seedlings were incubated for 48 h under static conditions in a plant chamber with a day/night cycle of 16 h light at 24°C and 8 h dark at 20°C. We conducted two mono-host evolution experiments, one using only ***<u>A</u>****. thaliana* (Bs_root_**<u>A</u>**) and another with **<u>T</u>**omato (Bs_root_**<u>T</u>**), as well as two host-cycling evolution experiments, which either started on ***<u>A</u>****. thaliana* (Bs_root_**<u>A</u>**T) or **<u>T</u>**omato (Bs_root_**<u>T</u>**A) and switched plant species after every third transfer. Therefore, four different evolution experiments each with five parallel lineages (A to E) were passaged on both *A. thaliana* and tomato seedlings for 20 transfers, spanning 40 days (**Fig. 1a** and **1b**). Consequently, every 2 days, a previously colonised plant seedling was transferred to a new sterile seedling, allowing only bacterial cells attached to the new root to continue in the experiment, enabling selection for a continuous cycle of dispersal, chemotaxis towards the plant root and biofilm formation on the root. To keep track and ensure sterile handling of seedlings during evolution, two controls without initial inoculant were included, in which either sterile seedlings or 10% of the MSNg medium were passaged. To follow the development of populations and test for contamination, bacterial cells colonising old roots, as well as the two controls, were plated on lysogeny broth (LB, Lennox; Carl Roth, Karlsruhe, Germany; 10 g/L tryptone, 5 g/L yeast extract, 5 g/L NaCl) agar medium after initial colonisation and after transfer 5, 10, 17 and 20.

**Fig. 1.**
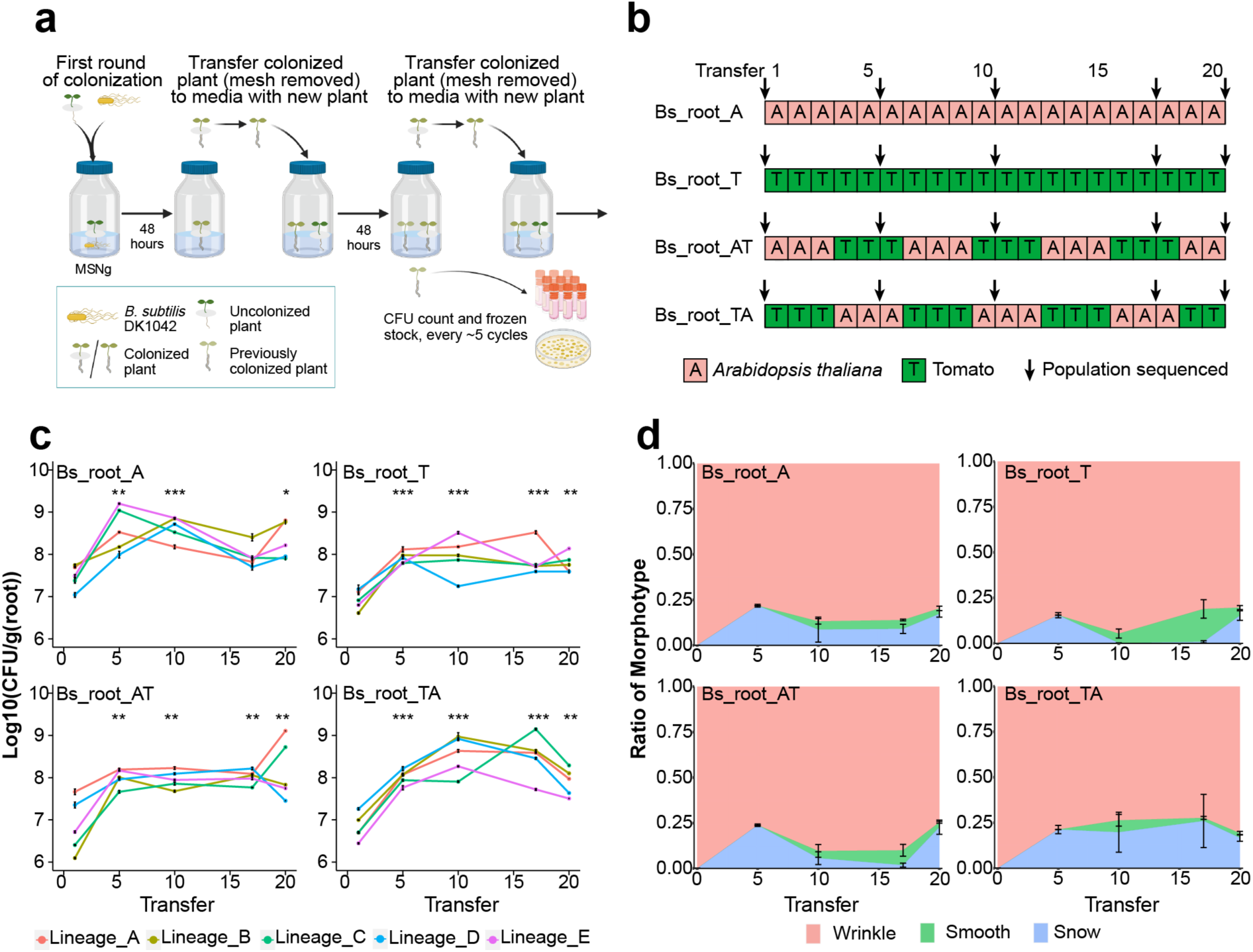
Evolution of *B. subtilis* on different plant roots. a. Experimental evolution scheme of *B. subtilis* adapting to plant seedlings (created with BioRender.com), **b.** Four evolution experiments were conducted, either as mono-evolution experiments with only *A. thaliana* or tomato seedlings, or cycling evolution experiments with plant species switched after every third transfer. Sequenced population samples are indicated by arrows. **c.** Root colonisation ability was measured as CFU per gram of root for T1, T5, T10, T17 and T20 in all lineages (n = 3, \**p* <0.05, \*\**p* <0.01, \*\*\**p* <0.001). **d.** Dynamic frequency changes of different morphotypes in the evolution experiment averaged for all five populations in each condition (n = 5). Results are means and error bars represent standard error in **c** and **d**.

### CFU quantification

In order to isolate biofilms from seedlings for quantification, roots were detached from sprouts, carefully washed in sterile MilliQ water and vortexed vigorously for 3 min in 1 ml of 0.9% NaCl in the presence of small glass beads (Ø0.75−1.00 mm; Carl Roth, Karlsruhe, Germany). The resulting cell suspensions were plated for colony-forming units (CFU) quantification and preserved as frozen stocks for subsequent comparison and DNA sequencing.

### Motility assays

A 20 mL volume of either 0.3% or 0.7% lysogeny broth (LB)-agar plates was used to test swimming and swarming motility. In both cases, plates were dried for 20 min before bacterial cultures were applied to the centre of plates. A 2 μL volume of overnight culture of ancestor or evolved isolates adjusted to an OD_600_ of 5.0 were spotted in the middle of the plate. To maintain similar humidity across all plates, multiple stacking was avoided. Plates were incubated at 30°C and images were taken for swimming (after 6, 8 and 24 h) and swarming (after 8, 10 and 24 h) using a Lumix DC T290 camera (Panasonic) equipped with a DC Vario-ELMAR lens (Leica). The area covered by bacterial cells at a given timepoint as well as the area of the whole agar plate were measured using Fiji ImageJ 1.52p. The capacity for swimming and swarming motility was calculated by dividing the covered area by the total area.

### Sequencing and variant calling

Archived samples from each experimental evolution environment were revived via cultivation in LB medium at 37℃ for 16 h. Genomic DNA was extracted from harvested cells of each culture using a EURx Bacterial and Yeast Genomic DNA Kit (EURx, Gdansk, Poland).

For whole-population genome sequencing of the evolved populations and ancestor strain samples, an MGIEasy PCR-free Library Prep Set (MGI Tech, Shenzhen, China) was used for library preparation, and 150 bp × 2 pair-end sequencing was performed on a DNBSEQ-Tx sequencer (MGI Tech) following the manufacturer’s procedures [28, 29]. More than 200× depth coverage of raw data was achieved for all population samples for further polymorphism calling.

To remove low-quality data, filtering was performed using SOAPnuke (version 1.5.6) [30]. Therefore, reads including >50% of bases with quality score <12, reads including >10% of unknown bases (N) and reads containing adaptor contamination were removed. Clean data were normalised to 200× depth for all sample to ensure similar variant calling sensitivity. Mutations were identified using *breseq* (version 0.35.7) with default parameters and a –p option for population samples [31, 32]. The default parameters reported mutations only if they included at least two reads from each strand and reached a frequency of at least 5% in the population. Mutations also present in ancestor strains were removed for further analysis.

### Genotype inferences from populations and Muller plots

To generate muller plots, the *Lolipop* (version 0.6) package (https://github.com/cdeitrick/lolipop) was used with default parameters [33, 34]. These methods predict genotypes and lineages based on the shared trajectories of mutations over time to test their probability of nonrandom genetic linkage. Muller plots were then manually coloured by the presence of shared mutated genes or genotypes. Genotypes and their frequencies were used for genotype diversity calculation in R (version 4.2.2) using the vegan (version 2.6-4) package.

### Jaccard Index calculation

Jaccard Index (*J*) for a pair of evolved lineages was calculated using formula (1). For a pair of lineages, each contains mutated gene set G1 and G2, respectively. In words, *J* is the number of genes mutated in both lineages divided by the total number of genes mutated in lineage 1 or lineage 2. *J* ranges from 0 to 1, with 1 indicating two lineages sharing exactly the same set of mutated genes and 0 indicating no mutated genes shared in the two lineages. This value was calculated for all pairs of evolved lineages, both within and between evolved environments.

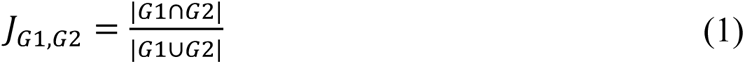

## Results

### *B. subtilis* showed rapid improvement and diversification during evolution on plant roots

We wondered whether plant-colonising bacteria adapt differently in an alternating host regime compared with a setup in which the same plant is used throughout the experiment. To investigate this, we explored adaptation of *B. subtilis* on the roots of two different plant species and compared the results with those of an experiment in which plant hosts were sequentially alternated. Evolved clones from the Bs_root_A setup have been examined in detail previously [27]. To evaluate the improvements in root colonisation during experimental evolution, CFU per gram of roots was measured after transfer 1, 5, 10, 17 and 20 (T1, T5, T10, T17 and T20, respectively). As expected [23, 24, 27], root colonisation ability improved significantly in all evolving lineages and conditions, by an average of 9.10-fold, 7.41-fold, 25.29-fold and 11.98– fold after 20 transfers in Bs_root_A, Bs_root_T, Bs_root_AT and Bs_root_TA, respectively (**Fig. 1c**). Furthermore, morphological diversification was also observed in all evolving lineages and conditions in CFU assays with morphotypes termed as Wrinkle (WR), Snow (SN) and Smooth (SM) [27], with colony morphology of the WR variant comparable to that of the ancestor. To investigate how the distribution of the distinct morphotypes developed during experimental evolution, overnight-grown cultures of frozen stocks of evolved populations from root 5, 10, 17 and 20 as well as from the ancestral strain (transfer 0) were prepared and plated on LB-agar. Different morphological variants were identified and counted to calculate the ratio of a given morphotype at each timepoint for each lineage. In all four conditions, WR was the most abundant in all lineages throughout the experiment (**Fig. 1d**). SN appeared in all lineages at T5, reaching an average frequency of 21.78%, 15.48%, 23.81% and 21.28% in Bs_root_A, Bs_root_T, Bs_root_AT and Bs_root_TA, respectively (**Fig. 1d**), but it was below the detection limit in some lineages at T10 and T17, then raised again in all lineages at T20. SM was observable in 11 of 20 lineages at T10 and only absent in one lineage at T17 and T20 (**Fig. 1d**). The three distinct morphotypes coexisted until the end of the evolution experiment, reaching average highly similar ratios of ∼80% WR, 17% SN and 3% SM in all four evolution experiments, revealing an astounding degree of parallelism at the morphotypic level.

#### Evolved isolates showed impaired motility

In a soil environment, bacterial cells must actively move towards plant roots during the initial phase of root colonisation, as well as when recolonising [21]. We hypothesised that, in the static hydroponic setup, which selects for regular root recolonisation including chemotaxis, biofilm formation and dispersal, motility would improve significantly during the evolution experiment, resulting in better root colonisation. To examine this, evolved end-point isolates from Bs_root_A and Bs_root_T were tested for swimming and swarming motility by spotting 2μL of an overnight culture on 0.3% and 0.7% LB-agar, respectively. Contrary to our expectations, none of the seven tested isolates evolved on *A. thaliana* or the five tested isolates evolved on tomato showed improvements in either swimming or swarming (**Fig. 2**). Instead, most isolates displayed a significantly smaller swimming radius after 6 h and 8 h compared with the ancestor, although all isolates colonised the entire plate after 24 h, thus mobility was preserved. Importantly, other forms of motility such as sliding [35, 36] might mask the effect of swimming after 24 h. Moreover, all isolates exhibited either delayed, reduced or even impaired swarming abilities. In future experiments, the frequencies of altered motility phenotype need to be assayed for each condition to clearly conclude selection against motility trait.

**Fig. 2.**
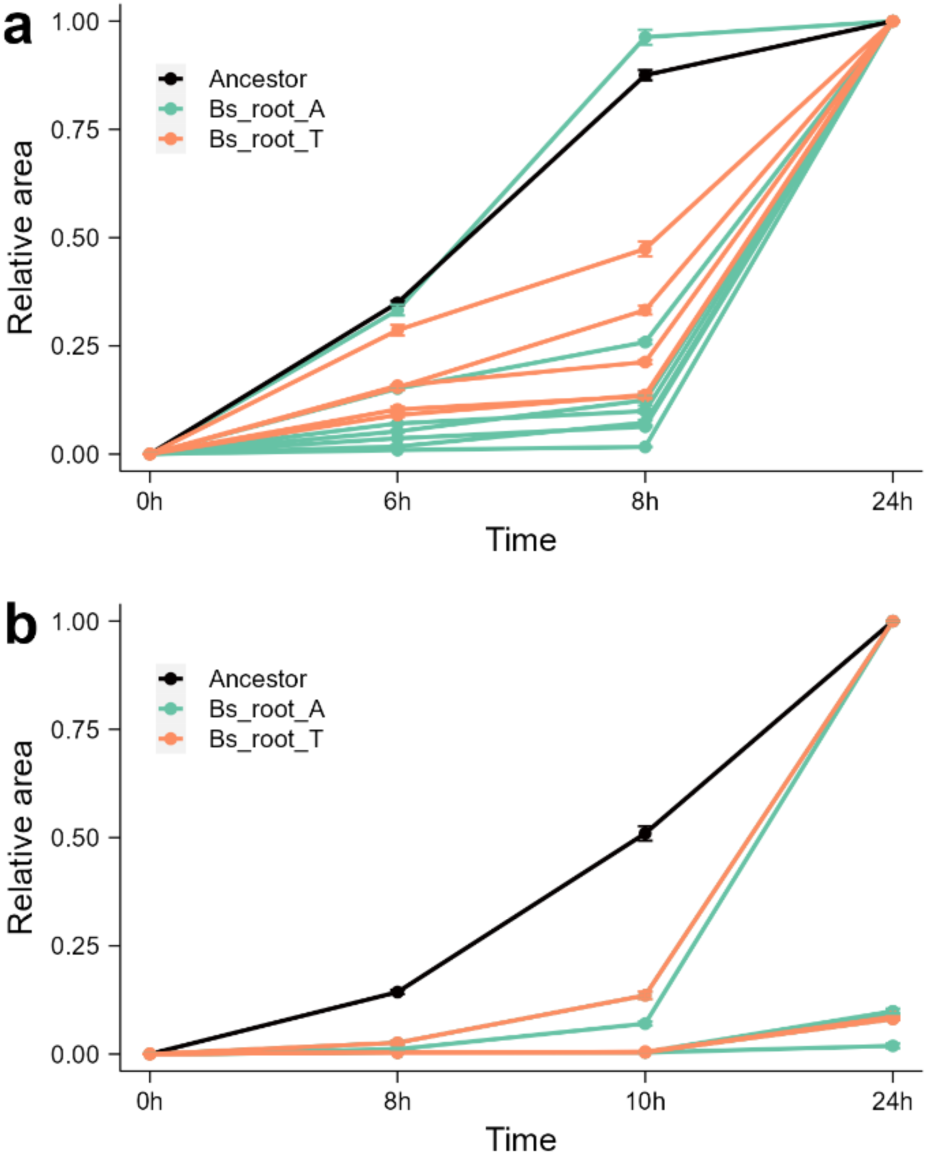
Evolved isolates showed impaired motility. Swimming. (a) and **Swarming (b)** motility of evolved isolates from Bs_root_A (lime green, seven isolates) and Bs_root_T (Soft orange, five isolates) compared with the ancestor (**Black**). Notably, swarming motility of several isolates were similar and low, and hence indistinguishable from each other in panel b. Results are means and error bars represent standard error, each isolate was tested in triplicate, and the ancestor was repeated 9 times. Tested isolates included two SN from lineage A and E of Bs_root_A, two SM from lineage A and E of Bs_root_A, three WR from lineage A, D and E of Bs_root_A, and five WR from lineage A (1), B (2) and C (2) of Bs_root_T.

#### Mutation spectrum and genetic diversity

Mutations provide the ultimate source of diversity, allowing selection to act and evolution to take place. To understand the genetic basis underlying the observed phenotypic diversification, impaired motility, and the dynamic landscape of evolution, we performed longitudinal whole-population genome sequencing of 79 *B. subtilis* population samples (one population sample was lost during freezer maintenance) and the ancestor strain, using the DNBSEQ-Tx platform [28, 29]. More than 200× coverage depth of raw data was achieved for each sample. Mutations, including single-nucleotide polymorphisms (SNPs) and short insertions and deletions (indels) were identified using the *breseq* pipeline (version 0.35.7) [31, 32]. The default parameters identified mutations only if they appeared in at least two reads from each strand and reached a frequency of at least 5% in the population. This analysis revealed 117, 106, 104 and 134 mutations in the evolved lineages from the Bs_root_A, Bs_root_T, Bs_root_AT and Bs_root_TA conditions, respectively (**Table 1**). No significant difference in mutation numbers were identified between the evolution conditions (Kruskal-Wallis test, *p* = 0.1277), though the mutation spectra were slightly different between the selection setups (**Table 1**). When the filtering step excluded mutations below 10% frequency, the cyclic-evolved conditions were found to contain more mutations than the mono-evolved conditions (*p* = 0.0005162, t = –4.2419, df = 17.542 via two-tailed t-test), suggesting that the mono-evolved conditions contained many low-frequency mutations, especially the Bs_root_A setup. Only four fixed mutations were present in all 20 lineages, possibly due to the short transfer times used in the experiment.

**Table 1.** Mutation type and number.

| Category | Bs_root_A | Bs_root_T | Bs_root_AT | Bs_root_TA |
| --- | --- | --- | --- | --- |
| Pseudogene | 0 | 1 | 0 | 0 |
| Non-coding gene* | 0 | 1 | 0 | 0 |
| Coding gene | 81 | 91 | 88 | 104 |
| Synonymous | 15 | 15 | 19 | 19 |
| Non-synonymous | 57 | 58 | 47 | 58 |
| Nonsense | 0 | 1 | 3 | 6 |
| InDel** | 9 | 17 | 19 | 21 |
| Intergenic*** | 36 | 13 | 16 | 30 |
| Number of mutations | 117 | 106 | 104 | 134 |
| Number of mutations (>10%) | 50 | 69 | 80 | 92 |
| Fixed mutations | 1 | 0 | 2 | 1 |

Note: There was no overlap between each mutation type. \* Non-coding genes include tRNA, ncRNA and repeat region (IS); \*\* InDel indicates small insertions or deletions in coding gene regions; \*\*\* Intergenic refers to all kinds of mutations occurring in intergenic regions.

Although there was no significant difference in the number of mutations between the evolution conditions, we predicted that populations evolved under cyclic conditions would show greater genetic diversity than mono-evolved populations at the end of the experiment due to the higher complexity and fluctuating nutrient composition caused by the alternating host plants. Therefore, the genotype diversities of endpoint samples were compared between conditions. Indeed, genotype diversity was significantly lower in mono-evolved conditions than in cyclic-evolved conditions (*p* = 0.01133, t = –2.9044, df = 14.315 via two-tailed t-test), and lowest in the Bs_root_A condition (**Fig. 3**; *p* = 0.1049, 0.03864 and 0.02767 via two-tailed t-test compared with Bs_root_T, Bs_root_AT and Bs_root_TA, respectively).

**Fig. 3.**
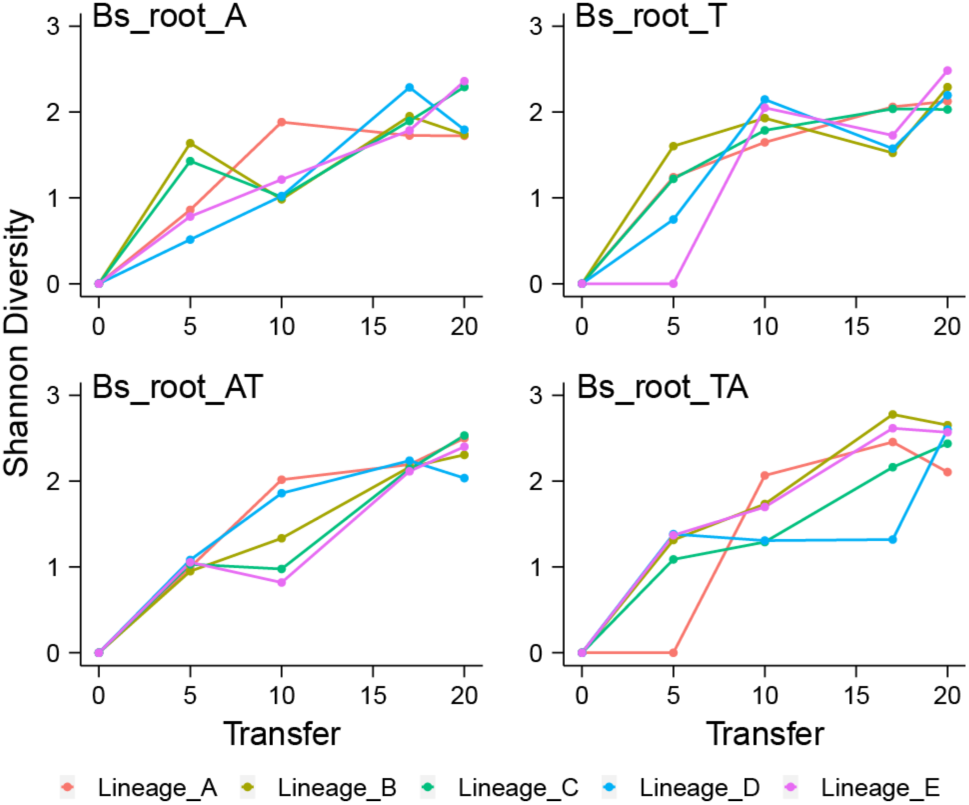
Genotype diversity. Dynamic distribution of genotype alpha diversity (Shannon method) in each population of four adaptation models over time. Genotypes and their frequencies are from genotype and genealogy analysis using *Lolipop*.

#### Ecological differentiation and parallel evolution

To infer the ecological differentiation in each lineage and visualise the changes in lineage frequencies from shared, nested mutation trajectories over time, we utilised the *Lolipop* software package developed by Cooper and colleagues [33, 34]. An average of 16 ± 1.10, 15.6 ± 2.15, 14.6 ± 1.85, and 17.4 ± 2.15 genotypes were obtained in Bs_root_A, Bs_root_T, Bs_root_AT, and Bs_root_TA conditions, respectively, with multiple mutations in each genotype. Consistent with the number of mutations, there were no significant differences in genotype numbers between evolved conditions (Kruskal-Wallis test, *p* = 0.3129). Overall, the number of genotypes continuously increased in all conditions, although it fell in some lineages at certain timepoints, consistently with the detected genotype diversity in each condition. Notably, neither genotype number nor diversity reached a peak within the time frame of the experiment (**Fig. 3**).

Each of the lineages was dominated by a few genotypes containing one or more mutations in certain genes. In general, *sinR*, *pstBA*, *yuxH*, *sigD* and eight flagellum-related genes were the most selected, demonstrating a high parallel evolution at gene and function levels within and between different evolved conditions (**Fig. 4**). Diverse functional genes were mutated, among which bacterial motility-related genes were targeted most frequently, indicating high parallelism at the functional level across all environments. To further compare genetic similarity between conditions, we determined the Jaccard Index (*J*) of all pairs of lineages within each condition. The Jaccard Index is commonly used for measuring similarity, indicating how likely it is for the same gene to be mutated in two separate lineages [37, 38], ranging from 0 to 1. Again, we observed relatively lower genetic similarity in mono-evolved conditions than in cyclic-evolved conditions, and lowest genetic similarity within the Bs_root_A condition, significantly lower than in Bs_root_T (*p* = 0.0249, t = –2.4471, df = 17.998 via two-tailed t-test) and the other two cyclic-evolved conditions (*p* <0.001 and <0.05, respectively; **Fig. 5**).

**Fig. 4.**
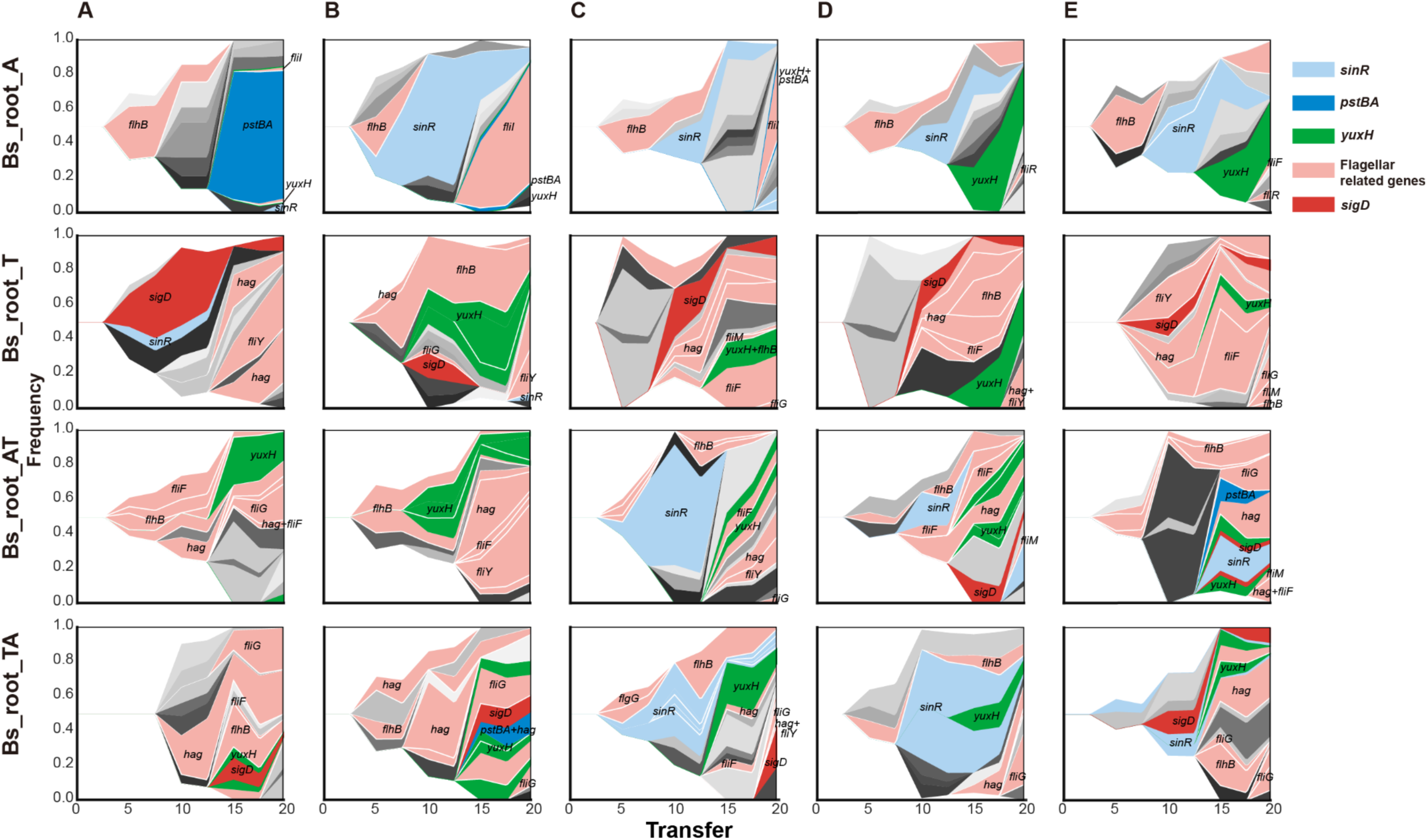
Evolutionary dynamics of *B. subtilis* during plant root colonisation. Each shade or colour represents a different genotype and vertical area corresponds to genotype frequency, inferred by *Lolipop*. Genotypes under selective pressure in certain conditions are highlighted in different colours and with annotations within or near shadows, while other genotypes are in different levels of grey without annotation. Nested genotypes of the highlighted genotype are highlighted with the same colour, except for nested genotypes which also contain genes under selective pressure. Notably, some genotypes are less abundant in the figure because their frequencies were similar to their nested genotypes (e.g., *fliI* and *yuxH* in lineage A of Bs_root_A, *pstBA* and *yuxH* in lineage B and C of Bs_root_A). Different colours represent different genotypes containing certain targeted genes, flagellar-related genes including *fliF*, *fliI*, *fliR*, *flhB*, *fliG*, *fliM*, *fliY*, and *hag*. Genotypes containing more than one targeted gene are coloured to match the gene displayed in the front.

**Fig. 5.**
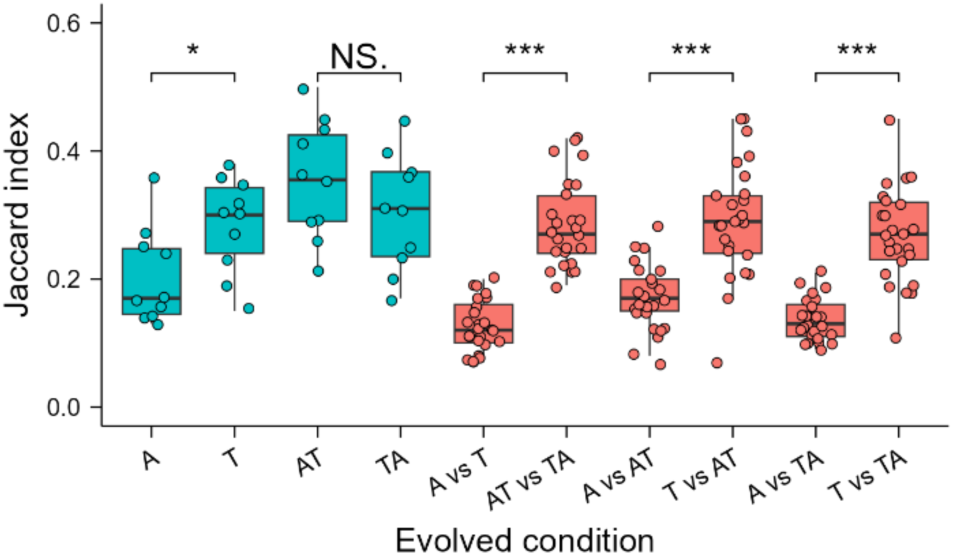
Parallelism. Degree of parallelism within and between each evolved environment was estimated using the Jaccard index. Asterisks indicate significant differences between different environment combinations (\**p* <0.05, \*\*\**p* <0.001, by student’s unpaired two-tailed t-test). Boxes indicate Q1−Q3, lines indicate the median, and circles indicate the *J* value of each pair within (blue) or between (pink) evolved conditions.

Subsequently, we compared the genetic similarity between different evolved conditions. The similarity between the two cyclic-evolved conditions was much higher than the two mono-evolved conditions (*p* = 2.628e-12, t = –10.022, df = 38.695 via two-tailed t-test), and the similarity between Bs_root_A and two cyclic conditions were also much lower than the similarity between Bs_root_T and two cyclic conditions (**Fig. 5**). These results strongly indicated that the selection pressure imposed by the *A. thaliana* was much weaker than observed with tomato, and the tomato root was potentially the main selective factor in the two cyclic-evolved conditions.

We observed high parallelism from root colonisation ability and morphotype to functional pathway and gene levels, as well as extensive parallelism at the level of individual nucleotides (**Dataset S1**). Strikingly, 18 of 20 lineages contained a frameshift in the *flhB* gene required for flagellum and nanotube assembly [39]. Another 8 bp deletion found in the *yuxH* gene, involved in c-di-GMP regulation in *B. subtilis* [40, 41], was detected in 15 lineages. Additional high parallelism mutations at the nucleotide level, present in more than 10 lineages and belonging to nonsynonymous mutations, occurred in *hag* and *sigD* genes. Notably, all mutations mentioned above reached a high frequency from 12.9% to 68.53% in the evolution conditions. We did not identify these mutations in the ancestor strain stock below a frequency of 5%, but we cannot rule out that they were already acquired in the overnight culture used to start the experimental evolution.

### *B. subtilis* adapts specifically towards different plant species

The lower mutation number (frequency >10%), lower genotype diversity, and lower genetic similarity of Bs_root_A strongly imply that the adaptation strategies between Bs_root_A and the other three conditions differed. Therefore, we focused on genes with a mutation frequency >10% and categorised them based on KEGG pathways. Overall, bacterial motility proteins, genetic information processing and metabolism-related genes were mutated more frequently and reached high frequencies, account for 61.36% of mutated genes (frequency >10%).

As mentioned above, bacterial motility proteins were the most selected in our experiment. Among the revealed motility protein genes, eight of eleven were flagellar structure protein-coding genes (**Fig. 6**) [42], and could be categorised into filament-coding genes (flagellin, *hag*), basal body-coding genes (*fliF*, *fliG*, *fliM*, and *fliY*) and flagellar type III secretion (T3S) apparatus protein genes (*fliI*, *fliR*, and *flhB*) [42]. Although bacterial motility genes were targeted frequently in all conditions, *fliI* and *fliR* were only mutated in Bs_root_A, and mutations in *fliG*, *fliY* and *hag* were only absent in Bs_root_A. FliI is a flagellar-specific ATPase functioning in the export of flagellar proteins [43], and FliR is part of the T3S apparatus [44] required for flagellum and nanotube assembly. Another gene required for flagellum and nanotube assembly, *flhB* [39], was mutated in all evolved conditions. FliG is involved directly in flagellar rotation[45], FliY interacts with the chemotaxis system and controls the direction of flagellar rotation [42, 46], and *hag* encodes the flagellin monomer protein Hag, which is also essential for flagellar assembly [47, 48]. The sigma factor-encoding gene *sigD*, which regulates flagella, motility, chemotaxis and autolysis in *B. subtilis* [48–50], was mutated in all except Bs_root_A samples, while *yuxH* (*pdeH*) encoding a phosphodiesterase that degrades c-di-GMP, was mutated in all conditions and relatively high frequencies. c-di-GMP is a second messenger that regulates diverse cellular processes in bacteria, including bacterial motility, biofilm formation, cell-cell signalling and host colonisation [40, 41]. Overall, species-specific and common mutations in both flagellar structural proteins and motility regulator proteins were identified, which were induced by species-specific evolution stress imposed by different plant species and the same static hydroponic setup.

**Fig. 6.**
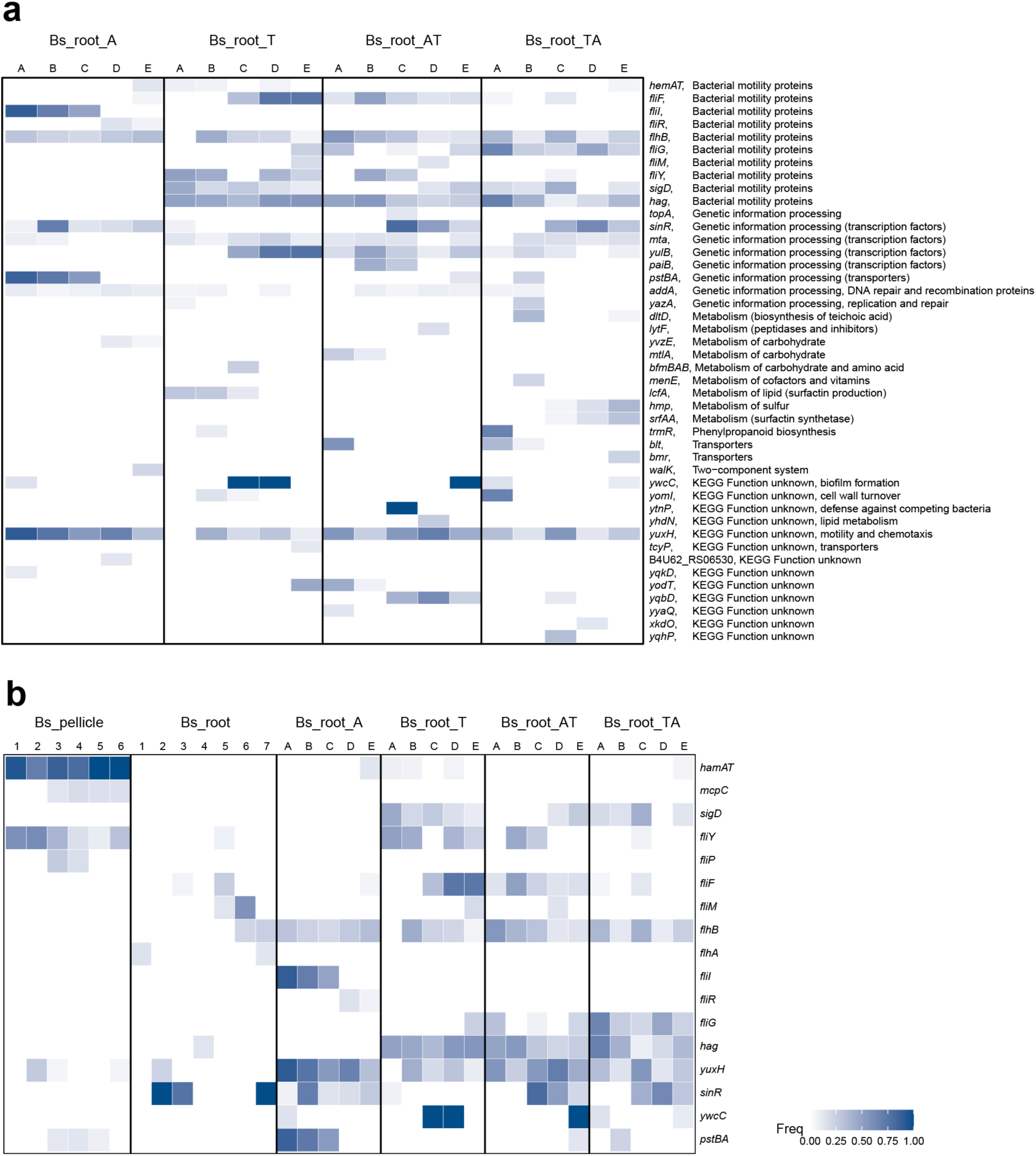
**Parallelism across different experimental setups. (a**) Genes with mutation frequency >10% in this study. (**b**) Mutated genes shared across different experimental setups. Each column represents one replicate lineage. In each lineage, the colour represents the highest frequency of mutations in that particular gene. Detailed information for these genes and mutations in **(a)** can be found in **Dataset S2**.

Mutations in biofilm development gene *sinR*, encoding a DNA-binding protein that regulates the expression of *epsA-O* and *tapA-sipW-tasA* operons involved in matrix production [51, 52], was only detected transiently with low frequency (maximum 7.19%) in Bs_root_T (**Fig. 4**, **Fig. 6** and **Dataset S2**), while it was observed more frequently and with higher frequency in the other conditions, suggesting an *A. thaliana*-specific mutation.

Another suspected *A. thaliana*-specific mutated gene, *pstBA*, encoding a phosphate transport system ATP-binding protein, was only absent in the Bs_root_T sample, and transiently with low frequency in the two cyclic-evolved conditions, but it was detected at high frequency in three of five lineages in Bs_root_A. The *pstBA* gene belongs to the five-gene *pst* operon, and it encodes a member of the PhoP-PhoR two-component signal transduction system [53–55] responsible for phosphate uptake under nutrient stress in *B. subtilis* [55], while the *pst* operon is alkali-inducible [53].

## DISCUSSION

We explored whether plant-colonising bacteria adapt differently in an alternating host regime compared with the same plant used throughout an experiment, and whether a bacterial species adapts differently to different plant hosts. To do this, experimental evolution and whole-population WGS sequencing were employed to analyse how *B. subtilis* evolves on *A. thaliana* and tomato seedlings, and when these two plant hosts were alternated in a static hydroponic setup [27]. The results did not provide a simple binary answer for the first question, and we observed species-specific adaptation patterns at the genetic level.

Similar to several previous studies [23, 24, 27], *B. subtilis* showed rapid improvement in root colonisation ability, albeit at varying rates, as three distinct morphotypes during adaptation in all evolved lineages and conditions. Additionally, the three distinct morphotypes coexisted until the end of the evolution experiment, reaching highly similar ratios in all four evolution experiments. More significantly, complementary effects were observed during root colonisation of *A. thaliana* seedlings through mixing of the three morphotypes from Bs_root_A [27]. Both root colonisation improvement and morphotypic diversification displayed a remarkable degree of parallelism across all evolved conditions. Furthermore, all but one randomly selected isolate from Bs_root_A and Bs_root_T showed impaired swimming and swarming motility. Populations from two host cyclic-evolved conditions shared numerous mutated genes with Bs_root_T, also suggesting similar trade-offs in the alternating plant setup. Analysis of the genetic basis underlying the two levels of parallelism (morphotypes and altered motility) revealed bacterial motility proteins, genetic information processing, and metabolism-related genes to be mutated more frequently and reaching high frequencies in all evolved conditions. Besides parallelism at phenotypic and functional levels, mutational resemblance was observed in certain genes, even at the individual nucleotide level. Two genes, *flhB* and *yuxH*, showed striking parallelism at both gene and individual nucleotide levels; 18 of 20 lineages contained a frameshift in the *flhB* gene and 15 lineages contained an 8 bp deletion in *yuxH*. The *flhB* gene is required for flagellum and nanotube assembly and *yuxH* is involved in c-di-GMP regulation in *B. subtilis* [40, 41], and both may contribute to adaptation of *B. subtilis* to plant roots by affecting motility. However, reintroduction of single mutations alone or in combination will be necessary in the future to precisely measure the influence of these mutations on fitness. Herein, we identified repeated changes across multiple levels of biological organisation from phenotypic, functional pathways and genes to individual nucleotides [37], demonstrating the overlapping evolutionary patterns employed by *B. subtilis* when adapting to *A. thaliana* and tomato roots, as well as roots in an alternating regime.

The static hydroponic setup used in our study selects for regular root recolonisation by bacterial cells on different plant species, including bacterial traits of chemotaxis, biofilm formation and dispersal [27]. Different histidine kinases influence the phosphorylation level of the global regulator Spo0A [15] when *B. subtilis* forms a biofilm on the roots of *A. thaliana* [16] and tomato [17], and different *Bacillus* species have distinct host plant colonisation abilities [19]. Based on previous results, we hypothesised that *B. subtilis* adaptation to *A. thaliana* or tomato will display distinct gene-level attributes underpinning species-specific adaptation. Although parallelism was identified at phenotypic and nucleotide change levels, obvious differences were detected in the number of mutations (at a frequency >10%), genotype diversity, mutated genes, and genetic similarity between and among different experimental conditions.

Fewer mutations (at a frequency >10%), lower genotype diversity and lower within-conditions genetic similarity were observed in mono-evolved conditions, especially in Bs_root_A. The difference between mono-evolved and cyclic-evolved conditions could be explained by a more heterogeneous, multi-resource, spatially-structured environment created by host-cycling, suggesting that environmental heterogeneity might have an important influence on adaptation [34, 37, 56]. When *Pseudomonas fluorescens* was evolved in five different environments varying in terms of resource type and arrangement, more mutations and higher degree of parallelism was observed at the gene level in the most heterogeneous environment [37]. Similarly, evolution of *Pseudomonas aeruginosa* populations on plastic beads that favour a biofilm lifestyle exhibited higher morphologic diversity and mutation rates than planktonic cultures [57]. Consistent with previous studies, the degree of parallel evolution was significantly higher in *B. subtilis* populations evolved in a heterogeneous, multi-resource, spatially-structured environment in the present work.

Bs_root_A populations harboured fewest mutations and the least similarity among these, even lower than Bs_root_T samples. Additionally, genetic similarity between the two mono-evolved conditions was much lower than between the two cyclic-evolved conditions, and similarity between Bs_root_A and the two cyclic conditions was also much lower than that between Bs_root_T and the two cyclic conditions. The lowest genetic similarity within Bs_root_A suggests that the fitness landscape in the *A. thaliana* evolving environment might have been more rugged than in tomato [58]. Furthermore, several specific mutated genes were also observed, which were present or absent exclusively in one plant species. These results imply that the adaptation patterns of *B. subtilis* to roots of *A. thaliana* and tomato differed, and *A. thaliana* exerted less influence on the adaptation process in the two cyclic-evolved conditions. A potential key difference between *A. thaliana* and tomato that might alter bacterial evolution is the composition and amount of exudates. In our experimental system, *B. subtilis* was evolved in minimal salt medium (MSM; see **Materials and Methods**), in which bacterial cells are dependent on carbon sources provided by the plant for their growth. The composition and amount of exudates in *A. thaliana* [59] and tomato [60] vary depending on growth stage.

We found that the most targeted genes were related to motility, while all lineages displayed improved root colonisation. Motility and biofilm formation are considered to be inversely regulated lifestyles during which bacteria express genes necessary for either motility or biofilm matrix production, but not both simultaneously [61, 62]. Motility is vital for many bacteria, as this process enables them to explore resources and supports the dispersal of progeny [63]. Motility is important for the effective root colonisation of different plant species by *B subtilis* under different conditions [18–21]. It is not surprising that the loss of motility is repeatedly evolved in populations growing in well-mixed liquid media where motility is not essential [37], and where the production of proteins for motility, primarily for flagella, is costly in terms of energy and resources [64]. A biofilm-motility trade-off has been suggested for *B. subtilis* evolved on *A. thaliana* roots under shaking conditions [24]. Surprisingly, all mutations in flagellar structure protein-coding genes were nonsynonymous SNPs or InDels, suggesting these mutations may cause loss-of-function of flagella. Notably, a 78 bp duplication (23 amino acids, residues 133−155) occurred in *hag* in three evolved conditions (12 lineages) and reached a high frequency, which would strongly affect the structure and function of flagellin, and therefore cell motility. Additionally, widespread and high-frequency mutations in *sinR* encoding a biofilm master regulator [65], *sigD* encoding sigma factor D [48–50] which regulates flagella, motility, chemotaxis and autolysis of *B. subtilis*, *yuxH* which is involved in c-di-GMP regulation in *B. subtilis* [40, 41], and *pstBA* which is responsible for phosphate uptake under nutrient stress in *B. subtilis* [55], were all predicted to regulate the motility-biofilm balance in *B. subtilis*, indicating that pleiotropy is widespread and can influence the adaptation of *B. subtilis*.

Finally, we compared the genetic similarities and differences in specific biological functions including aerotaxis, chemotaxis, motility (regulation) and biofilm development, when *B. subtilis* was evolved under different experimental conditions, including a static air-medium pellicle biofilm transfer mode (Bs_pellicle) [66], an *A. thaliana* root colonisation study under a shaking hydroponic setup (Bs_root) [24], and plant root colonisation under a static hydroponic setup (Bs_root_ATTA; **Fig. 6b**). Both common and condition-specific genes were found to carry mutations. We found that *hemAT* was mutated in both Bs_pellicle and Bs_root_ATTA systems, although less frequently in Bs_root_ATTA. Importantly, both experiments were performed under static conditions, where an oxygen gradient is generated in the culture, and HemAT is a key oxygen sensor important during pellicle formation [50]. The lower frequency of mutation in *hemAT* detected in Bs_root_ATTA suggests lower selection for aerotaxis in cells that colonise the submerged parts of plant roots compared with cells localised at the medium-air interface. Interestingly, the sigma factor-encoding gene *sigD* was mutated only in conditions with tomato serving as host in Bs_root_ATTA, suggesting that it is a tomato-specific mutational target gene. Flagella-related genes were found to be mutated in all three conditions, although mutations were detected in different genes and with different frequencies. Furthermore, the c-di-GMP-degrading phosphodiesterase-encoding *pdeH* was mutated in all three conditions, and more frequently in Bs_root_ATTA. The c-di-GMP second messenger modulates diverse cellular activities in bacteria including bacterial motility, biofilm formation, cell-cell signalling and host colonisation [40, 41]. However, previous studies on c-di-GMP signalling have mostly focused on Gram-negative bacteria. The *sinR* gene appears to be a target in plant-specific conditions because it lacked mutations only in Bs_pellicle samples. Another possible static condition-specific target similar to *hemAT* is *pstBA*, which is involved in high-affinity phosphate uptake in *B. subtilis* [55].

In conclusion, our study revealed parallel evolution across multiple levels of biological organisation from phenotypic to individual nucleotide levels, while the degree of parallel evolution at the gene level was significantly higher in populations evolved in a heterogeneous, multi-resource, spatially-structured environment. We also observed species-specific adaptation at the genetic level, possibly caused by selection stress imposed by different host plants. Additionally, a strong trade-off between motility and growth as well as biofilm formation was observed under the static selection conditions applied herein. Finally, we identified several condition-specific and common mutated genes by comparing genetic similarities and differences between three studies.

